# Defining traits of low-light adapted *Prochlorococcus* inhabiting surface waters of the Equatorial Pacific Ocean

**DOI:** 10.64898/2026.09.23.753768

**Authors:** Paul M. Berube, Trent LeMaster, Kelsy R. Cain, Yubin Raut, Zinka Bartolek, Shiri Graff van Creveld, Laure Arsenieff, Korinna Kunde, Sigitas Šulčius, Allison Coe, Nerissa L. Fisher, Stephen Blaskowski, Alexandra E. Jones-Kellett, Debbie Lindell, Randelle M. Bundy, E. Virginia Armbrust, Sallie W. Chisholm

## Abstract

A diverse array of photosynthetic phytoplankton drives primary production in equatorial surface waters. Among these, the cyanobacterium *Prochlorococcus* is an important contributor to net primary production in these typically iron-limited, high-nutrient and low-chlorophyll (HNLC) regions. Here, we explore the diversity of these organisms, in part, through targeted enrichment of *Prochlorococcus* cells using field-based high-speed cell sorting techniques. We demonstrate that the genomes of *Prochlorococcus* belonging to the low-light adapted LLI clade, and isolated from the surface of the Equatorial Pacific Ocean, are depleted in functions related to the assimilation of urea, nitrite, and amino acids. These are the first examples of LLI *Prochlorococcus* that have lost the ability to use nitrite, a trait considered to be a core feature of this clade. All new equatorial cultures of LLI *Prochlorococcus* appear to use a distinct isoform of protoporphyrinogen IX oxidase (HemG), for the biosynthesis of a chlorophyll precursor, that does not require the use of iron-containing heme. In contrast, the heme-dependent HemJ isoform is typically used by *Prochlorococcus* found outside equatorial HNLC waters. Together, these findings suggest that low-light adapted *Prochlorococcus* in the equatorial ocean possess accessory gene content that reflects adaptation to the generally iron-limited but nitrogen-replete conditions of surface waters.

## Introduction

The equatorial ocean is a dynamic environment where consistently high rates of marine primary production are observed (Le Bouteiller et al., 2003; Silsbe et al., 2016). Approximately 18% of global new production is estimated to occur in the Pacific equatorial upwelling system which is bounded by 5°N and 5°S (Chavez & Toggweiler, 1995; Le Bouteiller et al., 2003). Primary producers in this region are diverse, span both the Eukarya and Bacteria domains, and include a broad range of phytoplankton sizes (Landry & Kirchman, 2002; Landry et al., 2011; Le Bouteiller et al., 2003). This diversity is consistent with the broader observation that phytoplankton diversity tends to be higher in tropical regions and declines towards higher latitudes (Righetti et al., 2019).

Some regions of the equatorial Pacific are described as high-nitrate (or high-nutrient) but low in chlorophyll (HNLC), reflecting the concept that photosynthetic biomass is less than would be predicted given the relatively high concentrations of macronutrients such as nitrogen and phosphorus (Cullen, 1991). Instead of these macronutrients, iron availability is the factor that typically limits phytoplankton growth (Bonnet et al., 2008; Landry et al., 1997), including the cyanobacterium *Prochlorococcus* (Mann & Chisholm, 2000). *Prochlorococcus* is highly abundant in these equatorial regions with cell concentrations approximately an order of magnitude higher than the cyanobacterium *Synechococcus* and picoeukaryotic phytoplankton (Blanchot et al., 2001; Landry & Kirchman, 2002). Given the small size of *Prochlorococcus*, its contribution to gross primary production is estimated to be approximately 5-40% of the total in equatorial HNLC waters; measured contributions appear to depend at least partly on seasonal and decadal oscillations as well as the inherent dynamic variability of the equatorial ocean (Landry et al., 2011; Liu et al., 1997, 1999; Vaulot et al., 1995).

The diversity of *Prochlorococcus* encompasses a number of phylogenetic clades, many of which have distinct physiological adaptations and environmental distributions (Biller et al., 2015; Coleman & Chisholm, 2007; Moore et al., 1998; Rocap et al., 2002). These clades can be broadly grouped into a monophyletic high-light (HL) adapted cluster and a polyphyletic low-light (LL) adapted cluster (Biller et al., 2015). Among the high-light adapted *Prochlorococcus*, the HLI clade generally dominates in cooler oligotrophic waters, the HLII clade is broadly distributed across warm oligotrophic waters, and the

HLIII, HLIV, and HLV clades appear to be restricted to tropical ocean regimes limited by iron availability (Biller et al., 2015; Bouman et al., 2006; Johnson et al., 2006; Malmstrom et al., 2013; Rusch et al., 2010; Zinser et al., 2007). In contrast, clades within the highly diverse low-light adapted cluster of *Prochlorococcus* are generally restricted to the lower reaches of the euphotic zone — the LLI clade is an exception because it is tolerant of photon fluxes that are high enough to inhibit the growth of cells belonging to other low-light adapted clades (Biller et al., 2015; Malmstrom et al., 2010).

Among the *Prochlorococcus* that live in the surface waters of equatorial HNLC ocean regimes, cells belonging to high-light (HL) adapted clades have been observed to dominate. Strains belonging to the HLI and HLII clades have been isolated from surface waters of the Equatorial Pacific Ocean (Moore & Chisholm, 1999; Rocap et al., 2002) and partial single cell genomes are available for the HLIII and HLIV clades (Malmstrom et al., 2013; Pachiadaki et al., 2019). General features of adaptation by these clades to the iron-limited environments of tropical HNLC ocean regimes include the presence of siderophore transporters and the loss of several nitrogen assimilation pathways and Fe-containing proteins (Malmstrom et al., 2013; Rusch et al., 2010).

Although the Equatorial Pacific Ocean is a large and productive marine region with a high diversity of unicellular primary producers, there are relatively few isolates of *Prochlorococcus* from this environment. These cultivated representatives include several strains belonging to high-light adapted clades — MIT9215 (HLII clade), MIT9321 (HLII clade), MIT9322 (HLII clade), and MIT9515 (HLI clade) — as well as the low-light adapted MIT9211 strain which belongs to the LLII clade of *Prochlorococcus*. In order to further circumscribe the diversity and physiology of *Prochlorococcus* we aimed to isolate and characterize new strains from the equatorial HNLC surface waters.

We did so by employing high-speed cell sorting techniques and tailored nutrient amendments to enrich for *Prochlorococcus* cells uniquely adapted to life in the Pacific equatorial upwelling system. Although our primary targets were *Prochlorococcus* belonging to the uncultivated HLIII and HLIV clades, our efforts ultimately yielded new cultures of *Prochlorococcus* from the low-light adapted LLI clade.

## Materials and Methods

### Sampling locations

During two expeditions to the Equatorial Pacific Ocean — the SCOPE Gradients G4 cruise in November-December 2021 (TN397) and the SCOPE Gradients G5 cruise in January-February 2023 (TN412) — seawater was collected to initiate enrichment cultures for the isolation of picocyanobacteria. Both cruises repeated a portion of the 1992 U.S. JGOFS Equatorial Pacific Process Study along the 140°W line from north to south, passing through the Equator into the South Pacific Ocean. The Gradients G4 cruise was conducted during moderate La Niña conditions and the Gradients G5 cruise was conducted during a transition to El Niño conditions.

### Picocyanobacteria enrichment on Gradients G4

Seawater was collected from a trace metal clean diaphragm pump system deployed at a depth of 15 m at -2.5°N, -140°E, on 2021-12-06. The trace metal clean seawater was transferred to polycarbonate collection bottles (0.5 L) that were previously acid washed with 1 N hydrochloric acid, sanitized by microwaving for 10 minutes, and rinsed 3 times with the collected seawater. The seawater was then gravity filtered (i.e., no vacuum) through a 0.8 µm pore size and 47 mm diameter Whatman Nuclepore track-etched membrane filter (Cytiva Life Sciences, Wilmington, DE) to remove larger cells; the filtration units were previously acid washed with 1 N hydrochloric acid and sanitized by ultraviolet irradiation before the expedition. The filtrate was split into duplicate 200 mL aliquots. Each aliquot was amended with 16 µM ammonium chloride, 1 µM sodium phosphate, 0.090 µM manganese chloride, 0.008 µM zinc sulfate, 0.005 µM cobalt chloride, 0.003 µM sodium molybdate, 0.010 µM sodium selenite, 0.010 µM nickel chloride, and 11.7 µM sodium ethylenediaminetetraacetic acid. The ammonium chloride and sodium phosphate were treated with a Chelex-100 resin (Price et al., 1989) to remove trace metal contamination prior to use. One of the 200 mL aliquots was then amended with 0.1 nM iron chloride; at this total iron concentration, the free iron (Fe’) is predicted to be approximately 0.004 nM (Thompson et al., 2011). The other 200 mL aliquot was not amended with iron.

These aliquots (Fe-amended and Fe-unamended) were split into 8 mL picocyanobacterial enrichment replicates in 30 mL Oakridge polycarbonate tubes that had been acid-washed and microwave-sanitized. For the remaining 9 days of the expedition, these were incubated at 23.3-24.7°C and 8-11 µmol photons m^-2^ s^-1^ of white light provided by broad-spectrum LED lights. Onshore, the enrichment cultures were monitored by flow cytometry using a Guava easyCyte 12HT Flow Cytometer (Cytek BioSciences, Bethesda, MD) and transferred as needed in media prepared using microwave-sterilized seawater collected from the sample site using the trace metal clean pump system.

### High-speed cell sorting and picocyanobacteria enrichments on Gradients G5

On the Gradients 5 cruise, we physically selected *Prochlorococcus* cells from surface seawater samples using a shipboard Influx Cell Sorter (BD Biosciences) before amendment with nutrients. Seawater was collected from a trace metal clean diaphragm pump system deployed at a depth of 15 m at 0°N, -140°E, on 2023-02-06. Before sorting, the instrument was run using 18 megohm water as the sheath fluid, flushed with 0.2 µm filter-sterilized 10% bleach for 5 minutes, followed by 5 minutes of 0.2 µm filter-sterilized 18 megohm water, and then run dry before shutdown. The system was restarted with a new 0.22 µm Sterivex sheath filter and fresh 0.2 µm filter-sterilized saline sheath fluid (32 g/L of sodium chloride in 18 megohm water). The instrument was configured as follows: a 200 mW 488 nm laser that was run at full power during sorting; two 10% neutral density filters; small particle detection option enabled; data collection triggered by forward light scatter; and flow rate adjusted to ensure an event rate at *<*1000/sec to minimize particle interference and improve sample resolution. The sort settings were as follows: 86 µm tip size at 15 PSI; sort mode: 1 drop pure; piezo amplitude: 2.64 V; drop frequency: 36.58 kHz; and drop delay: 26.7167. A total of 220,509 - 600,000 *Prochlorococcus* cells — identified by their red fluorescence (692/40 nm) and forward scatter — were sorted into a 48 well plate with each well containing 500 µL of the original seawater collected using the trace metal clean pump system. Using a 48 well plate minimizes the flight time of the droplet through air compared to other collection vessels. The sorted cells were then transferred into a total volume of 10 mL of filter-sterilized seawater in an acid-washed and autoclave-sterilized 30 mL polycarbonate oakridge tube. These samples were amended with 20 µM ammonium chloride, 1 µM sodium phosphate, 0.090 µM manganese chloride, 0.008 µM zinc sulfate, 0.005 µM cobalt chloride, 0.003 µM sodium molybdate, 0.010 µM sodium selenite, 0.010 µM nickel chloride, 0.030 µM iron chloride, and 0.030 µM sodium ethylenediaminetetraacetic acid. The enrichments were incubated and monitored using the same methods used for the Gradients G4 enrichments as described above. Several of the successful enrichments were rendered axenic using dilution to extinction as previously described (Berube et al., 2015).

### Genome sequencing and assembly

Cultures (25 mL) were grown to mid-exponential phase, pelleted by centrifugation, resuspended in 50 µL of artificial seawater, and frozen at -80°C. DNA was extracted from the cell pellets using phenol/chloroform extraction (Wilson, 2001) that was modified with an additional final chloroform extraction step to remove any residual phenol. DNA was resuspended in TE buffer and DNA concentrations were determined using a PicoGreen dsDNA assay (Thermo Fisher Scientific, Waltham, MA). DNA sequence libraries were generated by the MIT BioMicro Center. PacBio HiFi (Circular Consensus Sequencing) libraries for the MIT2302, MIT2306, MIT2308, and MIT2309 cultures were sequenced at Harvard University’s Bauer Core Facility. High-quality Oxford Nanopore reads were generated for the MIT2304, G4PB16, G4PB17, and G5PB11 cultures at the MIT BioMicro Center. Reads were filtered using Filtlong version 0.3.1 (–min length 4kb –keep percent 85 –target bases 750mb) (Wick, 2025) and then used to assemble genomes with Flye version 2.9.6 (–genome-size 2m –meta) (Kolmogorov et al., 2019). Chromosomes were reoriented to start at the *dnaA* gene using dnaapler version 1.3.0 (Bouras et al., 2024) with default settings.

### Biogeography of LLI *Prochlorococcus* using 16S amplicon sequence variants

The Global rRNA Universal Metabarcoding Plankton (GRUMP) database (McNichol et al., 2025) was used to assess the distribution of LLI *Prochlorococcus* across the Pacific, Atlantic, and Indian Ocean basins. GRUMP ASVs that matched *Prochlorococcus* were reclassified at the clade level (Supplementary Materials and Methods). LLI ASV counts were normalized to the sum of the Cyanobacteria and chloroplast 16S counts, including those for ‘unclassified’ *Prochlorococcus* clades, which may include LLI sequences but were not classified as such due to the ASV matching multiple clades.

### Functional enrichment analysis

Contigs databases for the assembled genomes were generated and annotated using anvi’o version 9 (codename eunice) (Eren et al., 2021) — open reading frames were identified using pyrodigal (Camargo et al., 2024; Hyatt et al., 2010; Larralde, 2022) and then annotated using anvi-run-hmms, anvi-scan-trnas (Chan & Lowe, 2019), and anvi-run-ncbi-cogs with the COG24 database (Galperin et al., 2025). The predicted open reading frames were additionally annotated with the CyCOGv6 database by aligning the identified genes to CyCOGv6 proteins using blastp (-evalue 0.000001) (Camacho et al., 2009), and importing the best matching CyCOGv6 annotation (if any) into the anvi’o contigs database. A pangenome of LLI *Prochlorococcus* genomes (10 isolated from the Subtropical Pacific Ocean and 7 isolated from the Equatorial Pacific Ocean) was then generated using anvi-pan-genome (Delmont & Eren, 2018) in order to visualize the distribution of gene clusters across the genomes of LLI *Prochlorococcus* strains isolated from the Pacific Ocean. To avoid skewing the results of gene enrichment analysis and to mitigate the small number of equatorial genomes, three additional pangenomes were created using subsets of LLI genomes; in each of these pangenomes, all 7 newly isolated equatorial genomes were used alongside 7 randomly selected genomes isolated from subtropical waters. NCBI COGs (version 24) and CyCOGs (version 6) were used for functional enrichment using the anvi-compute-functional-enrichment-in-pan program (Shaiber et al., 2020) for each pangenome with 7 test genomes (equatorial isolates) and 7 control genomes (subtropical isolates).

### LLI Gene Frequencies in Pacific Ocean Metagenomes

The results of gene cluster enrichment analysis were validated by examining the frequency of NCBI COGs and CyCOGs in metagenomic data from the Pacific Ocean. Metagenomic sequence libraries were obtained from Tara Oceans (Pesant et al., 2015; Sunagawa et al., 2015), Bio-GO-SHIP (Larkin et al., 2021), and the Hawai’i Ocean Time-series (HOT) (Biller et al., 2018). We also produced sequencing data for the O2C-Floats (KM2418) cruise (Supplementary Materials and Methods, Supplementary Table S1). Paired-end reads were overlapped using bbmerge and then assigned to *Prochlorococcus* clades using the ProSynTax-workflow and the ProSynTax protein database (Coe et al., 2025). As part of this workflow, the number of genome equivalents of each clade was determined based on coverage of single-copy core genes. We then excluded metagenomes that had fewer than 5 genome equivalents of LLI *Prochlorococcus*. For the retained metagenomes (Supplementary Table S2), reads that mapped to LLI *Prochlorococcus* in the ProSynTax-workflow were extracted. These reads were annotated with NCBI COGs (version 24) and CyCOGs (version 6) using DIAMOND (version 2.1.11) blastx (Buchfink et al., 2021) with the NCBI COG24 protein database (Galperin et al., 2025) and the ProSynTax protein database (Coe et al., 2025) respectively. NCBI COG and CyCOG abundances in each metagenome were then determined by summing the length of amino acid residues aligned to each COG and then dividing by the average amino acid length of that COG. Frequencies per LLI genome were determined by normalizing NCBI COG and CyCOG abundances to LLI genome equivalents (Supplementary Tables S3 and S4). Differential NCBI COG or CyCOG representation in equatorial versus subtropical LLI genomes was assessed using the Mann-Whitney U test and the Benjamini-Hochberg correction for multiple hypotheses (Supplementary Tables S5, S6, S7, and S8), with the following COGs excluded from testing: (1) COGs where the mean frequency was 0 in both groups (i.e., absent in both equatorial and subtropical LLI genomes) or (2) COGs where the mean frequency was less than 0.2 in both groups (i.e., found in fewer than 1 genome at the minimum threshold of 5 LLI genome equivalents per metagenome library). All equatorial metagenomes examined were sampled from the upper 30 m of the water column while subtropical metagenomes were from the subsurface chlorophyll maximum layer where LLI cells typically dominate.

## Results and Discussion

### Isolation of novel *Prochlorococcus*

From trace metal clean seawater samples collected in the Equatorial Pacific Ocean, we ultimately generated 8 unialgal cultures of low-light adapted *Prochlorococcus* (Table 1) for which complete circular genomes were assembled from long-read sequences. Clade affiliations were determined using phylogenetic reconstruction based on shared single-copy core genes (Figure 1). One *Prochlorococcus* strain belongs to the recently described LLVIII clade (Becker et al., 2024), while the remaining 7 strains belong to the LLI clade (Table 1 and Figure 1). To the best of our knowledge, the only other low-light adapted strain to be isolated from this ocean region was the MIT9211 strain (LLII clade), but from a depth of 83 m (Kettler et al., 2007; Moore & Chisholm, 1999). Therefore, these are among the first LLI and LLVIII *Prochlorococcus* to be isolated from surface waters (15 m depth) of the equatorial ocean.

**Table 1.** Metadata for the isolated *Prochlorococcus* strains and unialgal enrichment cultures.

| Culture/<br>Strain ID | °N | °E | Collection<br>Date | Depth<br>(m) | Axenic<sup>a</sup> | Clade | Genus |
| --- | --- | --- | --- | --- | --- | --- | --- |
| G4PB16<sup>b</sup> | −2.5 | −140 | 2021-12-06 | 15 | No | LLI | *Prochlorococcus* |
| G4PB17<sup>c</sup> | −2.5 | −140 | 2021-12-06 | 15 | No | LLI | *Prochlorococcus* |
| G5PB11 | 0.0 | −140 | 2023-02-06 | 15 | No | LLI | *Prochlorococcus* |
| MIT2302 | 0.0 | −140 | 2023-02-06 | 15 | Yes | LLI | *Prochlorococcus* |
| MIT2304 | 0.0 | −140 | 2023-02-06 | 15 | Yes | LLI | *Prochlorococcus* |
| MIT2306 | 0.0 | −140 | 2023-02-06 | 15 | Yes | LLI | *Prochlorococcus* |
| MIT2308 | 0.0 | −140 | 2023-02-06 | 15 | Yes | LLI | *Prochlorococcus* |
| MIT2309 | 0.0 | −140 | 2023-02-06 | 15 | Yes | LLVIII | *Prochlorococcus* |

<sup>a</sup> Axenic strains were verified to be free of bacterial contamination by plating and flow cytometry.
<sup>b</sup> Low initial iron amendment for G4PB16.
<sup>c</sup> No iron amendment for G4PB17.

**Figure 1.**
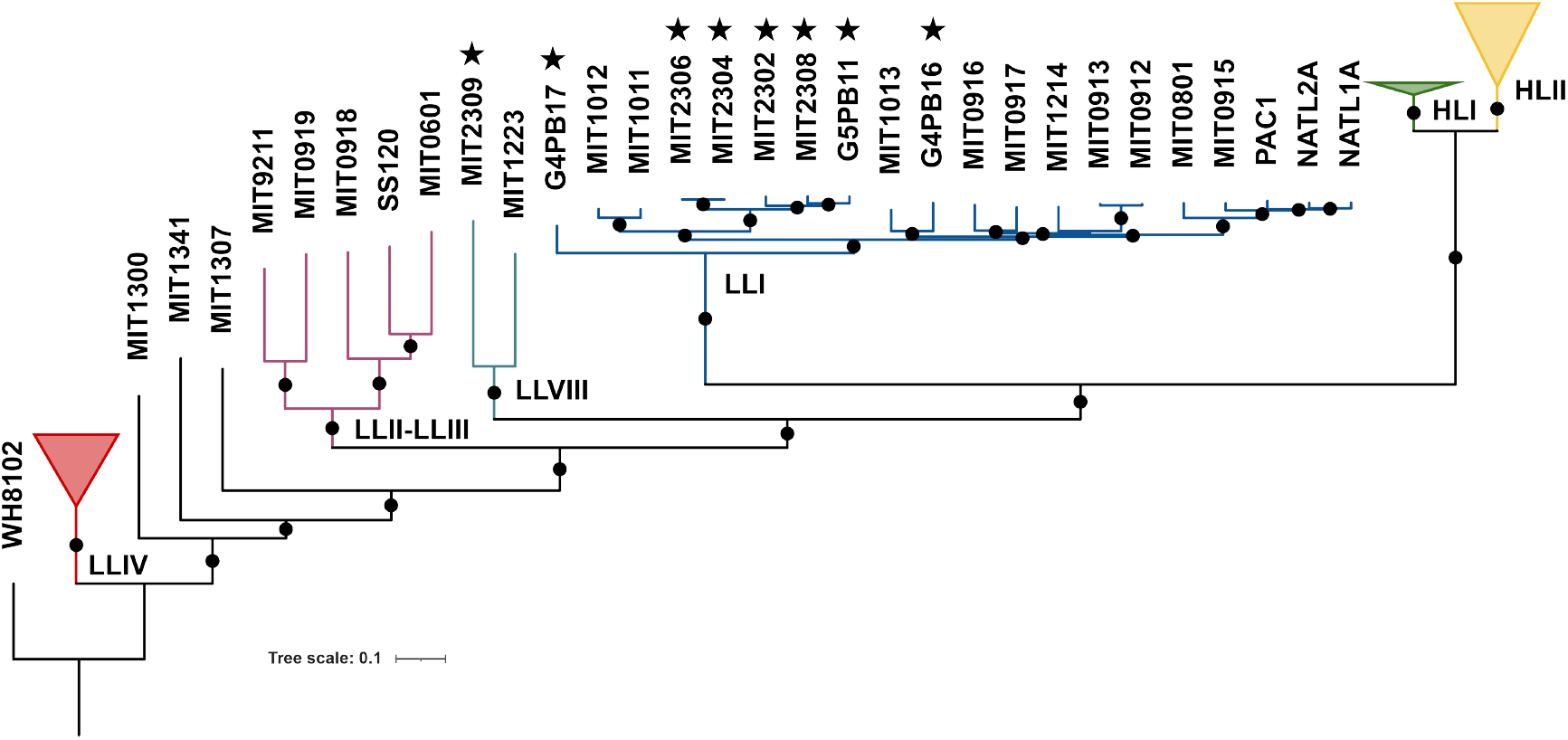
Phylogeny of *Prochlorococcus* isolates from the Equatorial Pacific Ocean. A phylogeny of new equatorial strains and representative *Prochlorococcus* cultivars — all with complete genomes — was reconstructed using an alignment of 424 single-copy core proteins. The tree is rooted on *Synechococcus* WH8102, which was included as an outgroup. Black circles indicate where the respective taxa are found on the same branch in *>*80% of 1000 replicate trees. Stars indicate the equatorial low-light adapted strains/cultures described in Table 1.

### Occurrence of LLI *Prochlorococcus* in tropical surface waters

Given that most of the new strains were from the LLI clade, we asked whether cells from this clade of *Prochlorococcus* were common in the surface waters of the Equatorial Pacific Ocean. In other words, do LLI *Prochlorococcus* rise to a high enough abundance in surface waters to contribute significantly to primary production? Or, did our enrichment methods simply select for cells that could best tolerate perturbations during enrichment and isolation? To address this, we turned to a large data set of amplicon sequence variants (ASVs) — the Global rRNA Universal Metabarcoding Plankton (GRUMP) database (McNichol et al., 2025). This data set is based on amplicons generated using universal rRNA gene primers, enabling the simultaneous profiling of the Bacteria, Archaea, and Eukarya domains (including chloroplast 16S rRNA gene sequences). Thus, in the context of assessing the contribution of LLI *Prochlorococcus* to the total surface phytoplankton community, this data set allows one to estimate the fraction of rRNA gene sequences — among Bacteria and Eukarya — that are LLI *Prochlorococcus*.

We observed that the fraction of LLI *Prochlorococcus* ASVs relative to the combined total of Cyanobacteria and chloroplast 16S ASVs in surface waters was elevated in regions that experience consistent vertical turbulence (e.g., the poleward edges of subtropical gyres and equatorial waters) relative to stratified gyre regions (Figure 2). This observation is consistent with previous work conducted at the Hawai’i Ocean Time-series in the North Pacific (22.75°N, -158°E) and the Bermuda Atlantic Time-series Study (BATS) in the North Atlantic (31.67°N, -64.17°E). At these subtropical locations, LLI *Prochlorococcus* were commonly observed in surface waters during winter mixing events while other low-light adapted clades were not (Malmstrom et al., 2010). Among low-light adapted *Prochlorococcus*, it was found that the LLI clade is unique in its ability to tolerate rapid increases in light intensity (Malmstrom et al., 2010), likely allowing this clade to persist when vertically mixed to the surface.

**Figure 2.**
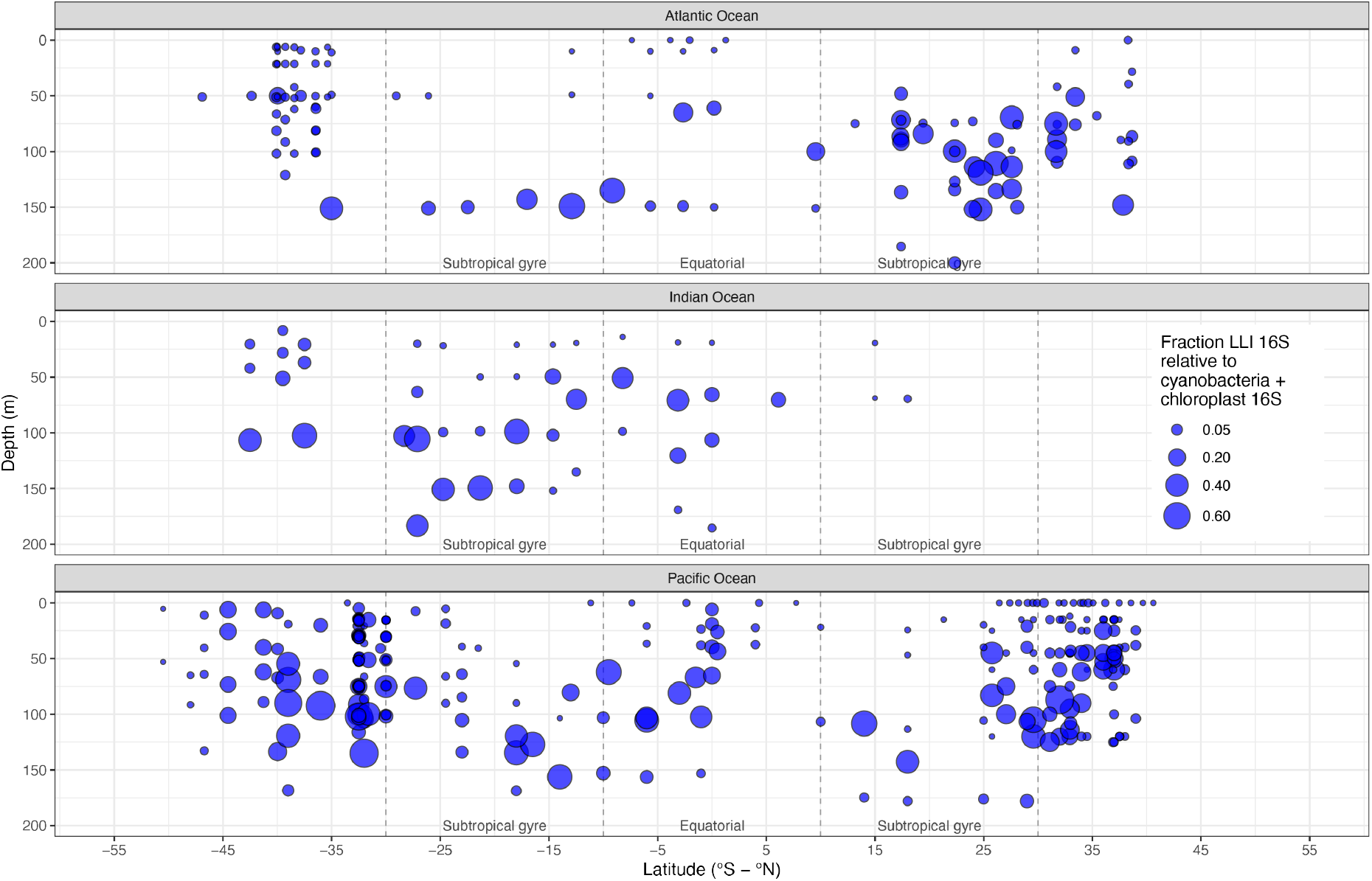
Global distribution of LLI *Prochlorococcus* ASVs relative to Cyanobacteria and chloroplast ASVs. Amplicon sequence variants (ASVs) in the GRUMP database (McNichol et al., 2025) that match the 16S rRNA gene of LLI *Prochlorococcus* relative to the total number of ASVs matching Cyanobacteria 16S rRNA and microalgal/eukaryotic phytoplankton chloroplastic 16S rRNA genes in the Atlantic, Indian, and Pacific Oceans. The size of each circle corresponds to the fraction of Cyanobacteria and chloroplast ASVs belonging to LLI *Prochlorococcus*.

Although seasonal signals are obscured in the GRUMP data set due to the absence of time-series data, the enhanced abundance of LLI *Prochlorococcus* in the top 50 m of the water column is particularly apparent for the Pacific Ocean (Figure 2); here, the GRUMP database includes samples collected approximately 10-30 degrees to the west of the location where our new strains of low-light adapted *Prochlorococcus* were derived. In these equatorial regions of the Pacific Ocean, while high-light adapted *Prochlorococcus* are indeed dominant (Supplementary Figure S1), LLI *Prochlorococcus* ASVs can reach 10-15% of all Cyanobacteria plus chloroplast ASVs in the top 50 m of the water column (Figure 2). These likely represent a lower bound of the proportional fraction of phytoplankton that are LLI *Prochlorococcus* because we required a unique match between an ASV and the LLI clade; further, chloroplasts of eukaryotes/microalgae often have 2 copies of the 16S rRNA gene while LLI *Prochlorococcus* only have 1. Many *Prochlorococcus* ASVs were unclassified (Supplementary Figures S2 and S3), some of which may be LLI *Prochlorococcus* that share a sequence variant with other clades. Given the diverse array of phytoplankton groups that coexist in the productive Equatorial Pacific Ocean, LLI *Prochlorococcus*, based on ASV data, may be a more important feature of this ecosystem than previously recognized.

### Pangenome of novel low-light adapted *Prochlorococcus* in tropical surface waters

We next assessed the gene content of our newly isolated strains of low-light adapted *Prochlorococcus*. Gene clusters that were identified in the 8 new low-light adapted *Prochlorococcus* strains were compared to previously sequenced isolates belonging to the same LLI and LLVIII clades of *Prochlorococcus* (Figure 3). Each new LLI genome from the equator possessed between 64 and 97 genes that were not previously observed in our CyCOGv6 database (Berube et al., 2018), while the equatorial LLVIII genome (MIT2309) had 139 genes that were previously unobserved. Many of these new gene clusters were unique to each new equatorial strain (Figure 3). Among these previously unobserved genes, all but 1-4 in each genome encoded hypothetical proteins, consistent with the vast majority of novel marine bacterial genes having an unknown function. Regardless, most gene clusters in each new genome could be assigned a CyCOG and many could be assigned a NCBI COG function (Figure 3).

**Figure 3.**
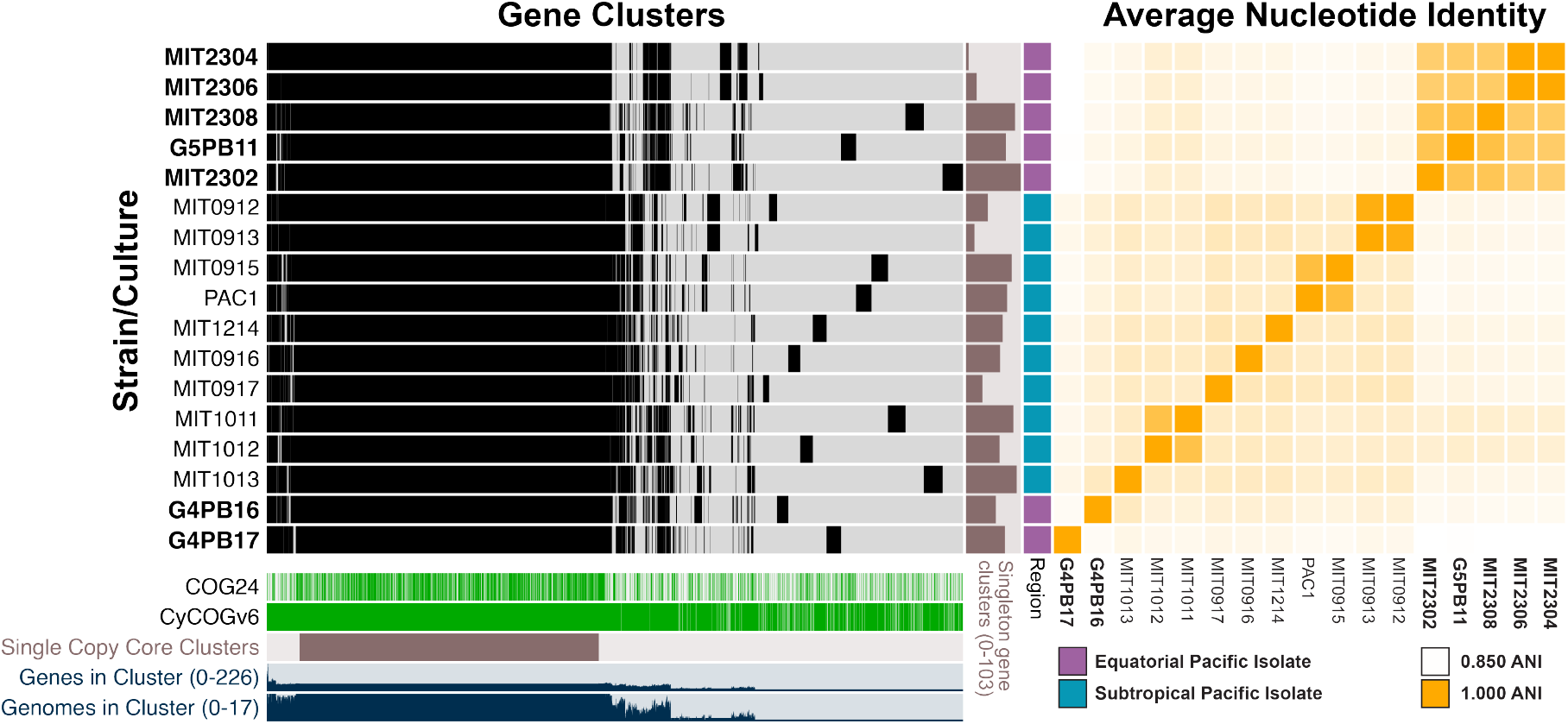
Pangenome of LLI *Prochlorococcus*. New LLI *Prochlorococcus* strains and enrichment cultures are shown in bold type. Gene clusters in each LLI *Prochlorococcus* genome are colored in black (present) or gray (absent). Below the presence/absence data are shown gene clusters annotated with an NCBI COG (COG24) or CyCOG (Berube et al., 2018), gene clusters observed as single-copy core gene clusters (present in all genomes), the number of genomes contributing to each gene cluster, and the number of genes in each gene cluster (normalized using a square root function to facilitate visualization). To the right of the presence/absence data are shown the number of singleton gene clusters (present in only one genome) in each strain and the location from which each strain was isolated (Equatorial Pacific, purple; Subtropical Pacific, cyan). Average nucleotide identity among strains is shown as a heat map (range 0.850-1.000). Ranges of values for bar plots are shown in parentheses.

### Genomic adaptations identified among equatorial LLI *Prochlorococcus* strains using functional gene enrichment

*Prochlorococcus* can be abundant in the surface waters of the equator (Figure 2); what adaptations do they possess that allow them to be successful in the dynamic, productive, high-nitrogen, yet generally iron-limited surface of the Equatorial Pacific Ocean? To address this question, we used gene enrichment analysis to identify genes under- or over-represented in the equatorial surface LLI genomes relative to LLI *Prochlorococcus* isolated from at or below the subsurface chlorophyll maximum layer of the subtropical gyres of the Pacific Ocean.

Given the high fraction of gene clusters annotated as a CyCOGv6 protein, we first looked at the distribution of CyCOGv6 functions among LLI *Prochlorococcus* (Table 2). We found that pathways for the transport and assimilation of urea and nitrite were absent in the new equatorial LLI *Prochlorococcus*. Notably, the trait for nitrite assimilation was thought to be a core feature of LLI *Prochlorococcus* based on an analysis of genomes derived from isolates and cultivation-independent single cells (Berube et al., 2019). Other CyCOGs were also observed less frequently among the equatorial LLI strains, although with false discovery rates that make their relevance more speculative. In particular, these included a nucleoside 2-deoxyribosyltransferase and a hypoxanthine phosphoribosyltransferase. These enzymes are involved in pyrimidine/purine recycling (Kilstrup et al., 2005), suggesting that this feature is more important in the typically nitrogen-limited subtropical gyres of the Pacific Ocean compared to the Pacific equatorial upwelling system.

**Table 2.** Proteins differentially encoded in *Prochlorococcus* from the Equatorial Pacific (7 genomes) relative to those from the Subtropical Pacific (10 genomes) based on Cyanobacterial Clusters of Orthologous Groups of Proteins Version 6 (CyCOGs).

| Product/Function | q-value Replicate<sup>a</sup> |  |  | CyCOG ID | Proportion<sup>b</sup> |  |
| --- | --- | --- | --- | --- | --- | --- |
|  | 1 | 2 | 3 |  | Gyre | Eq |
| Urea ABC transporter membrane protein | **0.035** | **0.031** | **0.035** | 60001300 | 1 | 0 |
| Urease accessory protein | **0.035** | **0.031** | **0.035** | 60001304 | 1 | 0 |
| Urease subunit gamma | **0.035** | **0.031** | **0.035** | 60001326 | 1 | 0 |
| Assimilatory nitrite reductase | **0.035** | **0.031** | **0.035** | 60001560 | 1 | 0 |
| Urease subunit beta | **0.035** | **0.031** | **0.035** | 60001317 | 1 | 0 |
| Urea-binding protein | **0.035** | **0.031** | **0.035** | 60001296 | 1 | 0 |
| Urea ABC transporter ATP-binding protein | **0.035** | **0.031** | **0.035** | 60001294 | 1 | 0 |
| Alpha-E superfamily protein | 0.134 | **0.031** | 0.147 | 60001386 | 0.9 | 0 |
| Nucleoside 2-deoxyribosyltransferase | 0.134 | 0.065 | 0.147 | 60001366 | 1 | 0.14 |
| Hypoxanthine phosphoribosyltransferase | 0.134 | 0.065 | 0.147 | 60001349 | 1 | 0.14 |

<sup>a</sup> Replicate gene enrichment analyses, each consisting of the 7 new equatorial strains and a random subset of 7 subtropical Pacific strains. q-values < 0.05 are highlighted in bold.
<sup>b</sup> The fraction of genomes encoding the CyCOG in each set of strains isolated from the North Pacific and South Pacific subtropical gyres (Gyre; N=10) and new strains isolated from the Equatorial Pacific (Eq; N=7).

The distribution of NCBI COGs among equatorial and subtropical LLI *Prochlorococcus* was generally consistent with the overall functions identified among the CyCOGs. Nitrogen assimilation functions — including transporters for the uptake of amino acids in addition to the transport and assimilation of urea — were depleted in the equatorial strains (Table 3). In addition, equatorial *Prochlorococcus* may potentially use an alternative isoform of protoporphyrinogen oxidase which is involved in the last shared step of heme and chlorophyll biosynthesis (Skotnicová et al., 2018). Most *Prochlorococcus*, overwhelmingly isolated from subtropical gyres, use the HemJ isoform that requires iron-containing heme as a co-factor. All of the newly isolated LLI *Prochlorococcus* from the equator appear to use the HemG isoform that uses iron-free flavin mononucleotide instead of heme. While statistical support for this difference is borderline due to the limited sample size of genomes (Table 3), we hypothesize that cells possessing a chlorophyll biosynthesis pathway that is less dependent on iron would be at a selective advantage in the iron-limited Equatorial Pacific Ocean.

**Table 3.** Proteins differentially encoded in *Prochlorococcus* from the Equatorial Pacific (7 genomes) relative to those from the Subtropical Pacific (10 genomes) based on NCBI Clusters of Orthologous Groups of Proteins (COGs).

| Product/Function | q-value Replicate<sup>a</sup> |  |  | COG ID | Proportion<sup>b</sup> |  |
| --- | --- | --- | --- | --- | --- | --- |
|  | 1 | 2 | 3 |  | Gyre | Eq |
| Branched-chain amino acid transport LivF | **0.021** | **0.019** | **0.020** | COG0410 | 1 | 0 |
| Branched-chain amino acid transport LivK | **0.021** | **0.019** | **0.020** | COG0683 | 1 | 0 |
| Branched-chain amino acid transport LivM | **0.021** | **0.019** | **0.020** | COG4177 | 1 | 0 |
| ABC-type transport system ATPase | **0.021** | **0.019** | **0.020** | COG4674 | 1 | 0 |
| Branched-chain amino acid transport LivH | **0.021** | **0.019** | **0.020** | COG0559 | 1 | 0 |
| Hydrogenase/urease maturation HypB | **0.021** | **0.019** | **0.020** | COG0378 | 1 | 0 |
| Urease accessory protein UreE | **0.021** | **0.019** | **0.020** | COG2371 | 1 | 0 |
| Urease accessory protein UreF | **0.021** | **0.019** | **0.020** | COG0830 | 1 | 0 |
| Urease accessory protein UreH | **0.021** | **0.019** | **0.020** | COG0829 | 1 | 0 |
| Urease beta subunit (UreB) | **0.021** | **0.019** | **0.020** | COG0832 | 1 | 0 |
| Urease gamma subunit (UreA) | **0.021** | **0.019** | **0.020** | COG0831 | 1 | 0 |
| Hypoxanthine phosphoribosyltransferase | 0.105 | 0.053 | 0.087 | COG2236 | 1 | 0.14 |
| Nucleoside 2-deoxyribosyltransferase | 0.105 | 0.053 | 0.087 | COG3613 | 1 | 0.14 |
| Alpha-E superfamily protein | 0.105 | **0.019** | 0.087 | COG2307 | 0.9 | 0 |
| Hydrogenase/urease accessory HupE | 0.252 | 0.053 | 0.087 | COG2370 | 0.8 | 0 |
| Protoporphyrinogen oxidase HemG | 0.252 | 0.053 | 0.087 | COG4635 | 0.2 | 1 |
| Protoporphyrinogen oxidase HemJ | 0.252 | 0.053 | 0.087 | COG1981 | 0.8 | 0 |

<sup>a</sup> Replicate gene enrichment analyses, each consisting of the 7 new equatorial strains and a random subset of 7 subtropical Pacific strains. q-values < 0.05 are highlighted in bold.
<sup>b</sup> The fraction of genomes encoding the COG in each set of strains isolated from the North Pacific and South Pacific subtropical gyres (Gyre; N=10) and new strains isolated from the Equatorial Pacific (Eq; N=7).

#### Distribution of genes in the wild

Methods used to isolate bacteria from natural systems have inherent biases, and thus, the resulting isolates may not reflect the characteristics of the dominant members of those systems. To verify our comparative genomics results, we turned to culture-independent metagenomic data to assess how well the gene cluster distributions among new strains mapped onto the frequencies of these genes in the wild. We first built a test group of 17 metagenomic sequence samples obtained from equatorial (-10°N to 10°N) regions of the Pacific Ocean (Figure 4 and Supplementary Table S2). Frequencies of CyCOGs and NCBI COGs among LLI *Prochlorococcus* in these metagenomes were then compared to a control group of 11 metagenomic sequence samples obtained from the subsurface chlorophyll maximum layer at HOT (Figure 4) in the North Pacific subtropical gyre.

**Figure 4.**
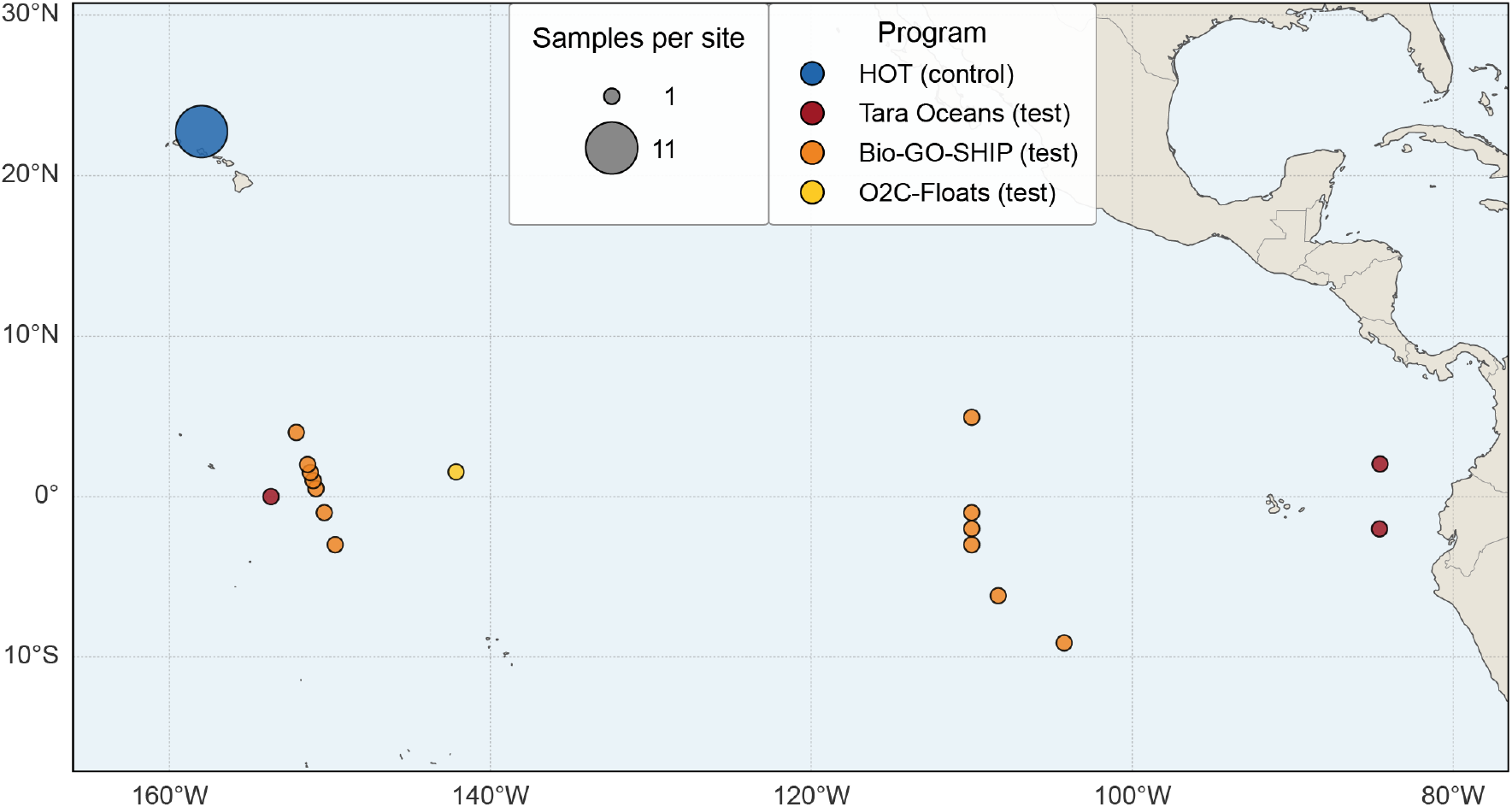
Locations of samples used for LLI gene enrichment analysis using metagenomic data. Eleven samples from the subsurface chlorophyll maximum layer at the Hawai’i Ocean Time-series were used as a control group (blue) alongside 17 samples collected in the Equatorial Pacific Ocean on Tara Oceans (red) and Bio-GO-SHIP (orange) cruises as well as the O2C-Floats cruise (yellow). The diameter of the circles is proportional to the number of metagenomic samples at the site.

The differential frequency of gene clusters observed in equatorial versus subtropical gyre metagenomes of LLI *Prochlorococcus* broadly recapitulated what we observed in the newly isolated equatorial strains — primarily, that genes encoding nitrogen uptake and assimilation proteins are depleted in LLI *Prochlorococcus* at the equator. Notably, the *nirA* assimilatory nitrite reductase gene (CyCOG60001560) was observed to occur in approximately 94% of LLI *Prochlorococcus* in the subsurface chlorphyll maximum layer of the subtropical gyres, but only in about 8% of LLI *Prochlorococcus* in the equatorial metagenomes (Supplementary Table S5). Additional nitrogen assimilation genes that were significantly depleted in the equatorial metagenomes included cyanate lyase (CyCOG60001979; COG1513) and the *focA* nitrite transporter gene (CyCOG60001803; COG2116) (Supplementary Tables S5 and S7).

The metagenomic data analysis also provided additional contextual information for functions that did not quite pass the 0.05 false discovery rate threshold in our comparative genomics analysis (Tables 2 and 3). For instance, while the genomic data suggested the potential for higher frequencies of nucleobase salvage functions among subtropical LLI *Prochlorococcus*, the comparative metagenomics analysis painted a more complicated picture. While genes encoding nucleoside 2-deoxyribosyltransferase and hypoxanthine phosphoribosyltransferase are indeed more frequent in the subtropical gyre metagenomes, genes encoding pyrimidine/purine nucleoside phosphorylase (PpnP) and aminopyrimidine aminohydrolase (TenA) were observed at higher frequency in equatorial metagenomes (Supplementary Tables S5 and S7). The former likely participates in purine salvage (Mao et al., 1997) while the latter is likely involved in the recycling of thiamine (Jenkins et al., 2007) in these *Prochlorococcus*. We also obtained further support that the HemG isoform of protoporphyrinogen oxidase, which uses flavin mononucleotide as a co-factor, is virtually absent from the subtropical gyre LLI *Prochlorococcus*. Cells in the gyres do appear to rely on the heme-dependent HemJ isoform, given that the *hemJ* gene was observed at a frequency of 1.37 *±*0.39 genes per genome equivalent in LLI *Prochlorococcus* in the North Pacific subtropical gyre (Supplementary Table S4). But, in contrast to our comparative genomics work, the metagenomics results suggest that equatorial LLI *Prochlorococcus* consist of cells that use a diversity of protoporphyrinogen oxidase isoforms, including the HemG isoform with a frequency of 0.34 *±*0.18 genes per LLI genome equivalent and the HemJ isoform with a frequency of 0.19 *±*0.16 genes per LLI genome equivalent (Supplementary Tables S4 and S7). We note that the sum of the HemG and HemJ frequencies is less than 1 for equatorial LLI genome equivalents; this may suggest the presence of divergent copies of these genes in these LLI *Prochlorococcus* (e.g., horizontally acquired from other organisms) or the use of alternative protoporphyrinogen oxidase isoforms that are not present in the CyCOGv6 database.

Equatorial LLI *Prochlorococcus* were also observed to possess some genes at a significantly higher frequency even though they were not found to be significantly enriched among the newly isolated equatorial strains. Beyond the alternative protoporphyrinogen oxidase, these include additional gene clusters that may also be selected for under iron-limiting conditions. Equatorial LLI *Prochlorococcus* appear to possess the FepB (COG0614), FepC (COG1120), and FepD (COG0609) components of an ABC-type siderophore transport system as well as the NfuA Fe-S cluster biogenesis protein (CyCOG60023405) — the latter of which is virtually absent in the LLI *Prochlorococcus* from subtropical gyres. Genes encoding the FepB, FepC, and FepD proteins were previously observed in high-light adapted genomes from the Equatorial Pacific Ocean (Malmstrom et al., 2013).

The new LLI *Prochlorococcus* strains from the Equatorial Pacific Ocean illuminate some of the adaptive features of a key primary producer in this dynamic, highly productive, and typically iron-limited marine environment. Although *Prochlorococcus* is known to be an abundant component of this environment, coexisting with *Synechococcus* and photosynthetic picoeukaryotes (Blanchot et al., 2001; Landry & Kirchman, 2002), the presence of low-light adapted clades in equatorial surface waters has been underappreciated. The LLI clade of *Prochlorococcus*, in particular, can rise to potentially more significant numbers in the surface waters of the Equatorial Pacific Ocean than previously recognized (Figure 2 and Supplementary Table S2). Similar to the uncultivated HLIII and HLIV clades of *Prochlorococcus* (Huang et al., 2012; Malmstrom et al., 2013; Rusch et al., 2010), it appears that LLI *Prochlorococcus* in HNLC regimes have dispensed with many nitrogen assimilation functions. This observation is consistent with a streamlined organism where selection operates to eliminate unnecessary functions (Díez et al., 2023; Giovannoni et al., 2014) in an environment where nitrogen is plentiful but iron is not. Still, the absence of urea uptake and assimilation genes in these new equatorial *Prochlorococcus* is intriguing, given that most *Prochlorococcus* have the genetic repertoire to use this nitrogen source and often prefer it (Berthelot et al., 2019; Díez et al., 2023). This suggests that equatorial LLI *Prochlorococcus* rely on ammonium as their primary source of nitrogen.

Equatorial LLI *Prochlorococcus* further exhibit adaptive features to life in an ecosystem where iron typically limits primary production — e.g., the presence of siderophore transport functions and the use of an alternative, heme-independent, protoporphyrinogen oxidase. Notably, the differential gene content that we observed was consistent with differential gene expression patterns found in metatranscriptomes (Blaskowski et al., 2025) from transects that crossed the North Pacific transition zone chlorophyll front (TZCF) (Polovina et al., 2017) — e.g., *Prochlorococcus* and *Synechococcus* urea and nitrite assimilation genes were highly expressed in the subtropical gyre and TZCF, but essentially undetected in subpolar HNLC waters.

Overall, our work has further circumscribed the diversity of an important marine primary producer through genomic characterization of newly isolated equatorial strains of LLI *Prochlorococcus* as well as comparative metagenomics of wild populations of an abundant low-light adapted clade of *Prochlorococcus*. Relative to LLI cells found at depth in the subtropical gyes, those at the equator exhibit defining adaptive features that likely allow them to persist in surface waters of the Pacific equatorial upwelling system.

## Acknowledgements

This work was supported by grants from the National Science Foundation (OCE-2048470 to P.M.B. and OCE-2220332 to F.H.F.), the Simons Foundation (Life Sciences Project Award ID 337262, S.W.C.; SCOPE Award ID 329108, S.W.C.; Award ID 723795, E.V.A.; Award ID 723552, K.K.; Award ID 721254, D.L.), and the Robert and Ardis James Foundation to S.W.C. The Simons Collaboration on Computational Biogeochemical Modeling of Marine Ecosystems supported Y.R. through Award ID 549931 to Michael J. Follows. We thank Dr. Angelicque White, Dr. Fernanda Henderikx-Freitas, and the captain and crew of the R/V Kilo Moana (KM2418) for their assistance with metagenomic sample collection, and Nhi Vo for assistance with submitting metagenomic data to NCBI. We thank the MIT BioMicro Center for assistance with DNA sequencing.

## Data and Code Availability

Assembled genomes and associated long-read sequencing data are available from the National Center for Biotechnology Information under BioProject PRJNA912210 and BioSample accession numbers SAMN61545810-SAMN61545817. Metagenomic data from the O2C-Floats cruise (KM2418) are available from the National Center for Biotechnology Information under BioProject PRJNA1244862. Code for running the workflows described herein are available at github.com/pmberube/ProSynTax-workflow-mergedreads and github.com/pmberube/equatorial-genomics-pipeline.

## Generative AI Disclosure

Claude Opus 4.7 and Claude Sonnet 4.6 were used to generate, debug, and edit analytic scripts and code; authors reviewed, tested, and validated all code. Claude Sonnet 4.6 was used to generate Python code to join and subset data and metadata tables; authors checked all results to confirm accuracy. Claude Sonnet 4.6 was used to refine LaTeX coding for manuscript formatting and Claude Opus 5 was additionally used to prepare and stage scripts and workflows for upload to the GitHub repository. No artificial intelligence tools were used for initial drafts or revisions of the manuscript. Claude Opus 5 was used to check the final manuscript for minor spelling and grammar errors. Authors take responsibility for the integrity of all content in this manuscript.

## Author Contributions

**Paul M. Berube**: Conceptualization, Data curation, Formal analysis, Funding acquisition, Investigation, Methodology, Project administration, Supervision, Validation, Visualization, Writing – original draft, Writing – review and editing; **Trent LeMaster**: Data curation, Investigation, Methodology, Validation, Visualization, Writing – review and editing; **Kelsy R. Cain**: Investigation, Methodology, Writing – review and editing; **Yubin Raut**: Formal analysis, Visualization, Writing – review and editing; **Zinka Bartolek**: Investigation, Writing – review and editing; **Shiri Graff van Creveld**: Investigation, Writing – review and editing; **Laure Arsenieff** : Investigation; **Korinna Kunde**: Conceptualization, Investigation; **Sigitas Šulčius**: Investigation, Writing – review and editing; **Allison Coe**: Investigation, Writing – review and editing; **Nerissa L. Fisher**: Investigation, Writing – review and editing; **Stephen Blaskowski**: Resources, Writing – review and editing; **Alexandra E. Jones-Kellett**: Resources, Writing – review and editing; **Debbie Lindell**: Conceptualization, Investigation, Funding acquisition, Writing – review and editing; **Randelle M. Bundy**: Conceptualization, Investigation, Funding acquisition, Resources, Writing – review and editing; **E. Virginia Armbrust**: Funding acquisition, Resources, Writing – review and editing; **Sallie W. Chisholm**: Funding acquisition, Supervision, Writing – review and editing

