## Supplementary Appendix for "Defining traits of low-light adapted *Prochlorococcus* inhabiting surface waters of the Equatorial Pacific Ocean"

#### Supplementary Materials and Methods

**Sample collection for metagenomics on the O2C-Floats cruise.** Replicate metagenomic samples were collected by filtering seawater onto 25 mm diameter 0.2 µm pore size Supor 200 membrane disc filters (Cat# 60301, Pall) during a transect from Hawai'i into the South Pacific Subtropical Gyre aboard the O2C-Floats cruise (KM2418) in October-November 2024 (Supplementary Table S1). Seawater (2-8 L per sample) was filtered using a peristaltic pump equipped with an inline filter. Following filtration, all samples were transferred directly into 2 mL cryovials, flash frozen using liquid nitrogen, and stored at -80°C until processing.

**DNA extraction and sequencing for O2C-Floats metagenomics samples.** The frozen samples were thawed briefly and then each filter was transferred from the 2 mL cryovial to a 2 mL polypropylene bead-beater tube. Immediately prior to DNA extraction, *Thermus thermophilus* genomic DNA (Cat# BAA-163D-5, ATCC) was added to the filters as an internal standard with a target of approximately 1% read abundance (Supplementary Table S1). Genomic DNA was then extracted from filters using a phenol/chloroform based method (Wilson, 2001) that was modified for *Prochlorococcus* as described by Coe et al. (Coe et al., 2025). Duplicate Illumina NexteraXT libraries (Cat# C-131-1096, Illumina, San Diego, CA) were prepared for each replicate DNA sample according to the manufacturer's guidelines for DNA templates. Samples were indexed using the TG Nextera XT Index Kit v2 (Cat# TG-131-2001, Illumina, San Diego, CA), with Set A indices used for the first replicate library and Set B indices used for the second replicate library. The exception was samples TL_01 and TL_02; for these, a single library was generated using Set A indices. Samples TL_01 and TL_02 were sequenced on the Illumina NextSeq 500 platform, while the remaining samples were sequenced on the NovaSeq 6000 platform at the MIT BioMicro Center. Paired-end sequencing reads were deposited in the NCBI Sequence Read Archive (Supplementary Table S1).

**Classifying *Prochlorococcus* ASVs by clade.** We developed an improved pipeline to annotate the clade for the majority of the 616 amplicon sequence variants (ASVs) labeled as *Prochlorococcus* in the Global rRNA Universal Metabarcoding Plankton (GRUMP) database (McNichol et al., 2025). First, we assembled a database of reference amplicons using the 1,106 *Prochlorococcus* genomes included in the ProSynTaxDB (Coe et al., 2025). 16S rRNA gene sequences were identified in each genome using Barrnap, imported into a QIIME 2 database (Bolyen et al., 2019), and trimmed using the universal primers 515Y and 926R that were used to amplify the sequences in the GRUMP database (McNichol et al., 2025). Incomplete sequences were removed from the database, resulting in 804 reference amplicons of 374 base pairs each with known clade identity. The clade of each *Prochlorococcus* ASV in the GRUMP database was then inferred by searching against the reference database using the 'classify-consensus-vsearch' function of the QIIME 2 'feature-classifier' module, using default settings and 'top-hits-only' set to true. This resulted in the successful classification of 586 of the ASVs. These were filtered in a final step to exclude clade-level classifications that were not unique (i.e., cases where an ASV had top hits to multiple clades); this yielded a final set of 451 ASVs in the GRUMP database with a predicted *Prochlorococcus* clade classification.

### Supplementary Tables and Figures

Supplementary Table S1. Metadata for O2C-Floats (KM2418) metagenomic samples.

Supplementary Table S2. Metagenomic samples used to examine differential frequencies of LLI gene clusters between equatorial and subtropical regions.

Supplementary Table S3. Frequencies of CyCOGs in LLI *Prochlorococcus* (genes per genome equivalent).

Supplementary Table S4. Frequencies of NCBI COGs (COG24) in LLI *Prochlorococcus* (genes per genome equivalent).

Supplementary Table S5. CyCOGs with differential frequencies between surface equatorial and subsurface subtropical samples; ordered by fold difference in the frequency of each CyCOG per genome equivalent.

Supplementary Table S6. CyCOGs found at similar frequencies between surface equatorial and subsurface subtropical samples.

Supplementary Table S7. NCBI COGs (COG24) with differential frequencies between surface equatorial and subsurface subtropical samples; ordered by fold difference in the frequency of each COG per genome equivalent.

Supplementary Table S8. NCBI COGs (COG24) found at similar frequencies between surface equatorial and subsurface subtropical samples.

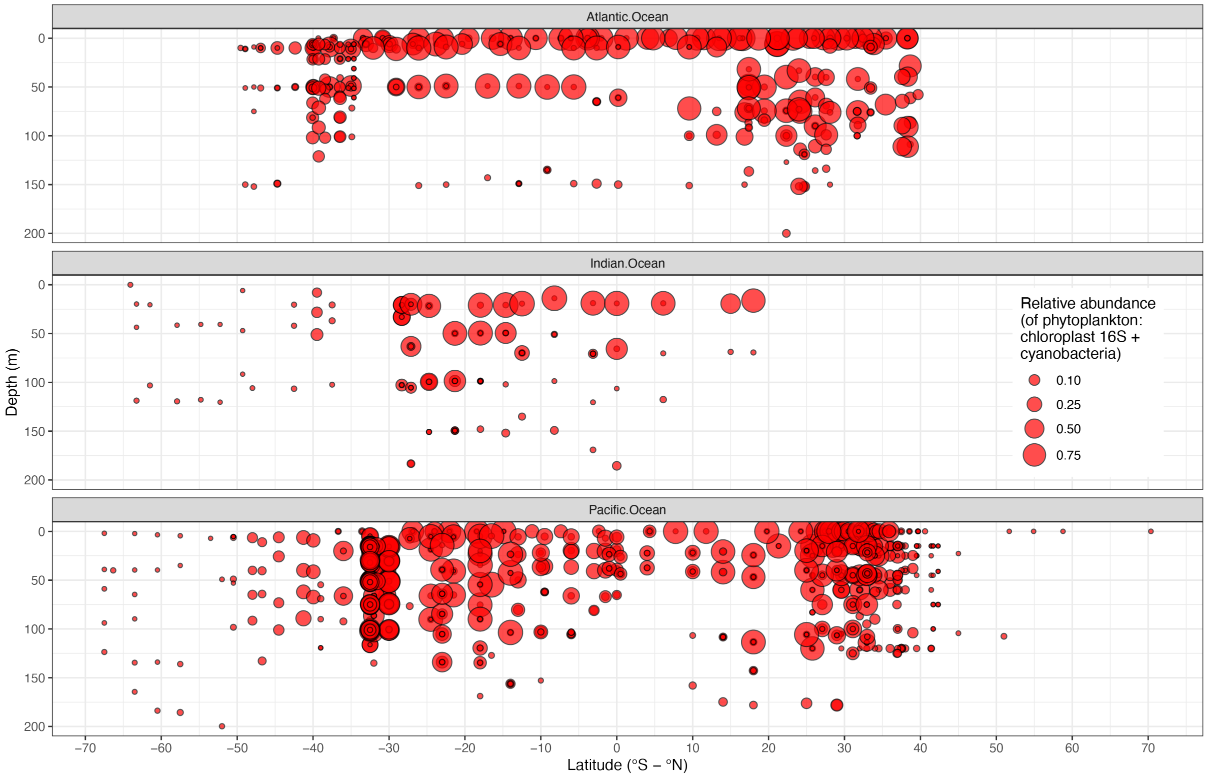


**Figure S1. Global distribution of high-light adapted *Prochlorococcus* ASVs relative to Cyanobacteria and Chloroplast ASVs.** Amplicon sequence variants (ASVs) in the GRUMP database (McNichol et al., 2025) that match the 16S rRNA gene of high-light adapted *Prochlorococcus* clades relative to the total number of ASVs matching Cyanobacteria and chloroplast 16S rRNA genes in the Atlantic, Indian, and Pacific Oceans. The size of each circle corresponds to the fraction of Cyanobacteria and chloroplast ASVs belonging to high-light adapted *Prochlorococcus*.

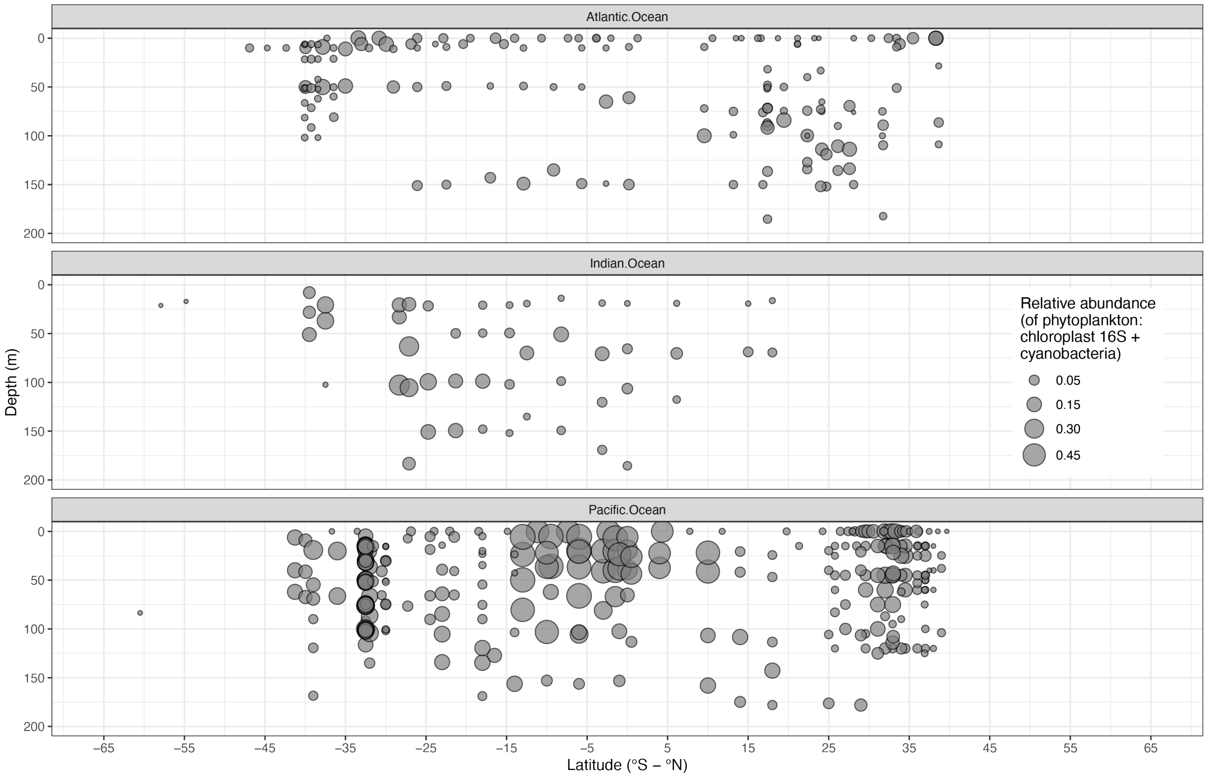


**Figure S2. Global distribution of Unclassified *Prochlorococcus* ASVs relative to Cyanobacteria and Chloroplast ASVs.** The relative abundance of unclassified *Prochlorococcus* ASVs (due to non-unique matches among clades) relative to the total number of ASVs matching Cyanobacteria and chloroplast 16S rRNA genes in the Atlantic, Indian, and Pacific Oceans. The size of each circle corresponds to the fraction of Cyanobacteria and chloroplast ASVs that were unclassified *Prochlorococcus*.

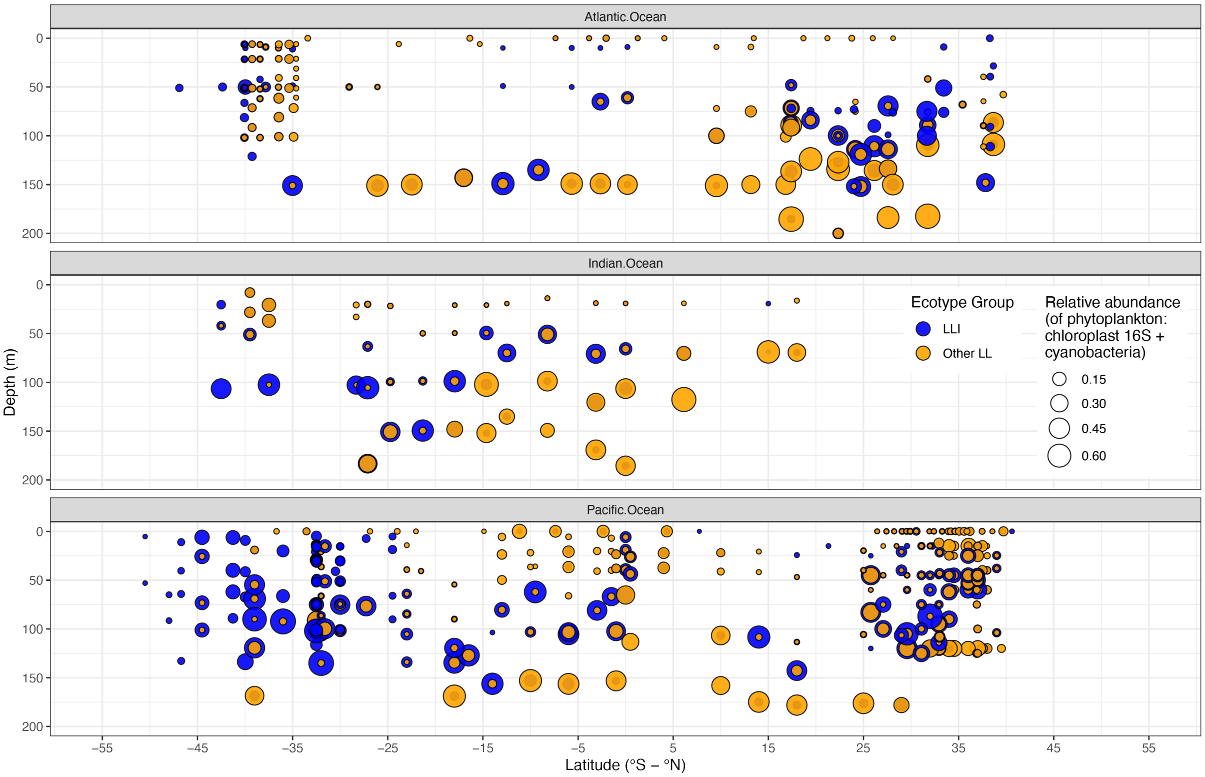


**Figure S3. Comparison of frequencies of LLI *Prochlorococcus* ASVs and Other LL *Prochlorococcus* ASVs relative to Cyanobacteria and Chloroplast ASVs.** Distributions of ASVs belonging to LLI *Prochlorococcus* and ASVs belonging to other low-light adapted clades (Other LL) are superimposed to visualize the relative contributions of these two *Prochlorococcus* subgroups in surface waters relative to deeper waters. The size of each circle corresponds to the fraction of Cyanobacteria and chloroplast ASVs belonging to these *Prochlorococcus* subgroups.

#### References


Bolyen, E., Rideout, J. R., Dillon, M. R., Bokulich, N. A., Abnet, C. C., Al-Ghalith, G. A., Alexander, H., Alm, E. J., Arumugam, M., Asnicar, F., Bai, Y., Bisanz, J. E., Bittinger, K., Brejnrod, A., Brislawn, C. J., Brown, C. T., Callahan, B. J., Caraballo-Rodríguez, A. M., Chase, J., Cope, E. K., Da Silva, R., Diener, C., Dorrestein, P. C., Douglas, G. M., Durall, D. M., Duvallet, C., Edwardson, C. F., Ernst, M., Estaki, M., Fouquier, J., Gauglitz, J. M., Gibbons, S. M., Gibson, D. L., Gonzalez, A., Gorlick, K., Guo, J., Hillmann, B., Holmes, S., Holste, H., Huttenhower, C., Huttley, G. A., Janssen, S., Jarmusch, A. K., Jiang, L., Kaehler, B. D., Kang, K. B., Keefe, C. R., Keim, P., Kelley, S. T., Knights, D., Koester, I., Kosciolek, T., Kreps, J., Langille, M. G. I., Lee, J., Ley, R., Liu, Y.-X., Loftfield, E., Lozupone, C., Maher, M., Marotz, C., Martin, B. D., McDonald, D., McIver, L. J., Melnik, A. V., Metcalf, J. L., Morgan, S. C., Morton, J. T., Naimey, A. T., Navas-Molina, J. A., Nothias, L. F., Orchanian, S. B., Pearson, T., Peoples, S. L., Petras, D., Preuss, M. L., Pruesse, E., Rasmussen, L. B., Rivers, A., Robeson, M. S., Rosenthal, P., Segata, N., Shaffer, M., Shiffer, A., Sinha, R., Song, S. J., Spear, J. R., Swafford, A. D., Thompson, L. R., Torres, P. J., Trinh, P., Tripathi, A., Turnbaugh, P. J., Ul-Hasan, S., van der Hooft, J. J. J., Vargas, F., Vázquez-Baeza, Y., Vogtmann, E., … Caporaso, J. G. (2019). Reproducible, interactive, scalable and extensible microbiome data science using QIIME 2. *Nature Biotechnology*, *37*(8), 852–857. https://doi.org/10.1038/s41587-019-0209-9

Coe, A., Mullet, J. I., Vo, N. N., Berube, P. M., Anjur-Dietrich, M. I., Salcedo, E., Parker, S. M., VonEmster, K., Bliem, C., Arellano, A. A., Castro, K. G., Becker, J. W., & Chisholm, S. W. (2025). A curated protein dataset for taxonomic classification of *prochlorococcus* and *synechococcus* in metagenomes. *Scientific Data*, *12*(1), 1895. https://doi.org/10.1038/s41597-025-06164-5

McNichol, J., Williams, N. L. R., Raut, Y., Carlson, C., Halewood, E. R., Turk-Kubo, K., Zehr, J. P., Rees, A. P., Tarran, G., Gradoville, M. R., Wietz, M., Bienhold, C., Metfies, K., Torres-Valdés, S., Mock, T., Eggers, S. L., Jeffrey, W., Moss, J., Berube, P., Biller, S., Bodrossy, L., Van De Kamp, J., Brown, M., Sow, S. L. S., Armbrust, E. V., & Fuhrman, J. (2025). Characterizing organisms from three domains of life with universal primers from throughout the global ocean. *Scientific Data*, *12*(1), 1078. https://doi.org/10.1038/s41597-025-05423-9

Wilson, K. (2001). Preparation of genomic DNA from bacteria. *Current Protocols in Molecular Biology*, *56*(1), 2.4.1–2.4.5. https://doi.org/10.1002/0471142727.mb0204s56
